# Adaptation to a host-associated lifestyle drives convergent loss of flagellar motility in Pseudomonadota

**DOI:** 10.1101/2025.11.22.689265

**Authors:** Roberto Siani, Caroline Gutjahr, Michael Schloter

## Abstract

**Background:** Host-associated bacteria must balance the benefits of motility through flagella against the offsets of energetic costs and immune surveillance. Understanding the interplay of evolutionary forces shaping this complex trait can provide insights on the dynamics and extent of within-host adaptations of flagellar assembly. We compared prevalence, redundancy, and homology of 55 known flagellar assembly genes across genomes of free-living and host-associated bacteria from a collection covering the entire Pseudomonadota phylum.

**Results:** We show that the isolation of host-associated bacterial populations leads to the erosion of the flagellar regulon, driving widespread flagellum loss across lineages. This gene loss is rarely compensated by horizontal gene transfer. Moreover, host association imposes a diversifying selective pressure that acts unevenly across pathway components, resulting in greater functional heterogeneity than in free-living bacteria.

**Conclusions:** Our results provide valuable insights into the distribution of flagellar genes in the phylum Pseudomonadota and its relation to bacterial lifestyle.

## Introduction

Bacteria loose, re-purpose, and acquire novel genes, in remarkably short periods of time when adapting to novel hosts [1–3]. In host-associated bacteria, mutations through genetic drift may become fixed quickly due to isolation [4]. The accumulation of neutral and deleterious mutations [5] can lead to loss or diversification of genes and, over time, degradation of the genome [6]. Survival within a host also requires adaptation to host control mechanisms (for instance, the immune system), which are hypothesized to be a driver of evolutionary changes within the microbiome [7]. One bacterial trait important for host-association is the flagellum [8]. While primarily acting as a propeller for motility, flagella also participate in mechanosensing, adhesion, chemotaxis, and facilitate complex behaviours such as colonization and biofilm formation [9–11]. Multiple studies have explored the origins of the bacterial flagellum. An early study identified a conserved set of flagellar genes, suggesting the flagellum arose through duplication and diversification of a single sequence [12]. Later phylogenetic analysis of flagellar genes challenged this view, proposing diverse, independent origins for flagellar genes and highlighting the role of horizontal gene transfer in their spread across bacterial phyla [13].

Whether from simple or complex origins, the bacterial flagellum is an evolutionary ancient and carefully constructed nano-machine: the reference assembly pathway maintained by the KEGGontology includes 55 orthologs [14] with structural and regulatory roles. The pathway, which was firstly mapped in *Escherichia coli* and *Salmonella typhimurium*, is organized in hierarchical layers and regulatory loops that orchestrate the timely expression of its genes [15]. Briefly, the master regulator, FlhDC (or species-specific alternatives such as FlrA and FleQ), initiate σ^70^ (RpoD) or σ^54^ (RpoN) dependant transcription of tier II operons. These operons encode components necessary for the assembly of the flagellar type III secretion system (fT3SS), the M/S ring, the switch complex, and the MS-rod junction. Once the fT3SS is assembled, the anti-sigma factor FlgM is exported, allowing FlhDC, together with σ^70^/σ^54^ and σ^28^ (the flagellar-specific sigma factor FliA) to drive the transcription of mixed tier II and III operons. These operons encode structural components of the basal-body and hook. Finally, σ^28^-dependant transcription of tier III operons completes the process, leading to the assembly of the motor/stator and the filament (flagellin proteins, primarily FliC).

Despite the advantage of flagella for locomotion and biofilm formation, the assembly, operation and maintenance of flagella demand significant energy resources [16], compromising growth rates in nutrient-poor environments [17]. Besides, as flagella are associated with virulence in several pathogenic species [18–23], animal and plants have evolved specialized receptors to recognize the main constituent of the flagellum, the filament protein FliC, as a marker of potential pathogens [24–26]. Thus, in addition to the advantage of saving energy, counter-adaptations must balance the trade-off between retaining functional flagella while evading recognition by the host immune system [27, 28]. Extensive research has been conducted on the polymorphism of FliC, which play a potential role in immune evasion [29–31].

Apart from *fliC,* a broader pattern of flagellar gene mutations has been observed in various host-associated clades [32–34], to different effects. For instance, a study on convergent adaptations across 29 human pathogenic species identified the flagellar regulon as a prominent mutational hotspot [35], except for the filament protein, which was not mutated in all species. In fact, fine-tuning of flagellar function has been shown to predominantly arise from mutations at regulatory checkpoints, rather than at structural components [36]. On the other extreme, flagellar motility is entirely lost in aphid endosymbionts, for whom the process of degeneration likely started with loss of the regulatory genes *flhD* and *flhC* [37] and may have involved the repurposing of flagellar basal bodies into secretion systems [38, 39].

While adaptation to specific hosts is known to impact *fliC* and flagellar motility, how host-microbe interactions influence the full complement of flagellar genes is not understood. We conducted a phylogenomic analysis of Pseudomonadota and investigated whether the rates of gene loss differ between host-associated and free-living bacteria, are homogeneous across the regulon, and driven by selective pressure. To disentangle the contributions of selection and neutral genetic drift, we account for genome erosion as a factor influencing evolutionary trajectories. Additionally, we examine the homology of flagellar orthologs to assess the role of diversifying selection in host-associated bacteria.

## Method Details

### Pseudomonadota genome collection

1839 complete genomes of representatives Pseudomonadota (previously known as Proteobacteria) were retrieved from NCBI RefSeq [40] archives (date of accession: 14.10.2023) using NCBI datasets v14.1 [41]. Sequences from the Alpha-, Beta-, Gamma-proteobacteria were included as less than 10 genomes were available for the other Pseudomonadota classes.

Sequences were decorated with one of two association states (host-associated, free-living), based on combined metadata from NCBI and BacDive [42]. Contextual information deposited on NCBI was prioritized. When information on the isolation source was absent or conflicting, BacDive was consulted. Coding and translated amino-acid sequences were predicted de-novo with Prodigal v2.6.3 [43].

### Phylogenomic tree reconstruction

A single-copy gene tree was constructed using GToTree v1.8.4 [44] with the available hidden Markov model (HMM) set for Pseudomonadota, gene-hits exclusion criteria of 0.33 and no filtering of genomes. Briefly, HMMER v3.2.2 [45] was used to identify 119 single-copy genes. Genes shorter than 0.67 x median length of the gene set or longer than 1.33 x median length of the gene set were considered spurious hits and removed. An individual alignment is produced for each gene set using muscle v5.1 [46]. The alignments are trimmed with trimal v1.4. rev15 [47] and concatenated. Phylogenetic estimation of the tree is conducted with FastTree2 v2.1.11 [48]. Protein alignments of the single-copy core genes were provided as input. Tree inference used BLOSUM45 amino-acid distances, balanced joining, and 1000 SH-like support replicates. The search strategy employed the default NNI and SPR optimizations (two SPR rounds with a radius of 10) and per-iteration ML-NNI optimization. FastTree’s default “TopHits” heuristics were used (1.00×√N, refresh threshold 0.80). Model fitting applied the Jones–Taylor–Thornton (JTT) substitution model with CAT rate heterogeneity using 20 rate categories.

### Identification of flagellar genes

GToTree optionally accepts a list of targets KEGG [49] orthologs (KO) families. Fifty-five KOs of orthologs belonging to the flagellar assembly pathway (flagellar) KO02040 were supplied as targets. GToTree sources profile HMMs for the targets and uses KOfamScan [50] to detect them in the genome collection, resulting in a count of number of gene copies per genome. KEGG profile HMMs uses family-specific adaptive score threshold to control false positive rates.

For homology modelling, a methodology proposed by Wheeler and colleagues [51] was adapted. HMM log-odds bitscores capture goodness of fit of queries and targets and are provided as a metric of the quality of the match. Given that HMM models emphasize conserved regions in their scoring, differences in bitscores (delta-bitscore) between sequences were used as a measure of conservation. New HMM profile was generated by aligning the KOfamScan detected sequences to their relative KOfam HMM profile. The newly generated profiles were used to screen the collection using hmmer v3.4.

### Phylogenetic analyses

All statistical analyses were conducted using R Statistical Software v4.3.2 [52] in RStudio v2023.12.0+369 [53], with the extensive use of tidyverse [54]. Firstly, the single-copy gene tree was coerced to ultrametric by extending all external edges using phytools v2.1-1 [55].

Multichotomies were resolved using the package ape v5.7.1 [56]. The tree was rooted at midpoint using phangorn v2.11-1 [57] and used for modelling phylogenetic correlation structures.

Prevalence and number of copies per gene, as detected by KOfamScan, were modelled using phylogenetic generalized (logistic and Poisson family) linear models from the package phylolm v2.6.2 [58]. For the bitscore modelling, a single best-hit per protein was retained for each genome. Bitscores were standardized and fit to a phylogenetic linear regression model from the package phylolm. In all three models, the logarithmic length of the genome was included as a covariate. For both the prevalence logistic model and the bitscore linear model, 999 independent bootstrap replicates were used to estimate 95% confidence intervals.

## Results

### Distribution of essential and accessory flagellar genes

To test whether the rate flagellar genes loss differs between host-associated bacteria and free-living ones, we collected 1839 genomes including 599 Alpha-, 318 Beta-, and 922 Gamma-proteobacteria, spanning 597 genera of Pseudomonadota. We classified 952 (52%) and 887 (48%) genomes as isolated from environmental (free-living bacteria) and host-derived materials (host-associated bacteria), resulting in balanced groups of free-living and host-associated bacterial genomes (Supplmentary Data 1). Our collection-wide search of the 55 flagellar genes composing the reference KEGG flagellar assembly pathway retrieved matches for 53 of them, spread across the whole phylogeny. We did not find any homolog of 2 genes, coding for the flagellar biosynthesis fusion protein FliR/FlhB (K13820) and the flagellar motility protein FlgQ (K24346) which is described only in *Campylobacter jejuni [59]*. Among putatively motile bacteria, i.e. those with at least one copy of the flagellin-coding gene *fliC* (n = 1234, [60]), we identified three groups of genes that consistently, frequently, or rarely co-occur with *fliC* (Supplementary Data 2). Out of the genes retrieved in our collection, 14 occur in less than half of the *fliC*-positive bacteria (rare flagellar genes). Master regulators of the assembly pathway were found at surprisingly low prevalence (34% for *flhDC*, 36% for *flrA/fleQ/flaK*). Other rare genes have regulatory functions (*flhE*, *flrC*, *fliZY*) or encode components of the H-ring and proteins associated to the outer membrane (*flgOPT*) and motor (*motCD*, *motXY*). Thirteen genes were found at moderate prevalence (accessory flagellar genes), between 50% and 80% of the *fliC*-positive bacterial genomes. These include the gene coding for the flagella-specific sigma-factor (*fliA*) and anti-sigma factor (*flgM*), for filament chaperones and cap (*fliDST*), for hook-filament junction, hook-length control and assembly chaperon (*fliK, flgL, flgN*), and for rod (*flgF*) proteins, the fT3SS ATPase, associated proteins (*fliHIJ*) and putative regulator/chaperone (*fliO/fliZ*). Finally, 25 protein-coding genes constituted a high prevalence cluster (core flagellar genes). This last group included genes coding for non-flagellar specific sigma factors (*rpoN*, *rpoD*), most structural components and regulators of the fT3SS (*fliPQR*, *flhAB*), of the switch complex (*fliLMN*, *fliFG*), and basal body/hook (*flgA*, *flgBCDEGHIJK*, *fliE*), and the stator proteins (*motAB*).

### Host association drive loss of flagellar genes

For both free-living and host-associated bacteria, larger genomes retained more detectable flagellar genes, with a significant association for 40 out of 53 genes. Host-association resulted in significantly lower prevalence of 12 of the 53 tested flagellar genes (*flgN*, *flgHIL*, *flhA*, *flhC*, *fliDS*, *fliHK*, *fliR, fliQ*), irrespective of genome size (Fig. 1, Supplementary Data 3). We found that only *fliY*, and the σ^54^ factor coding gene *rpoN* were more conserved in host-associated than free-living bacteria. In addition, we modelled dependence of number of gene copies on host-association using a phylogenetic Poisson regression (Supplementary Fig. 1, Supplementary Data 4). We observed that *fliC*, *fliL*, *motA* and *motB*, were often present in more than one copy (respectively, 53%, 49%, 60% and 53% of the times). For these four orthologs, our model predicted a significant and negative effect of host-association on number of copies.

**Figure 1.**
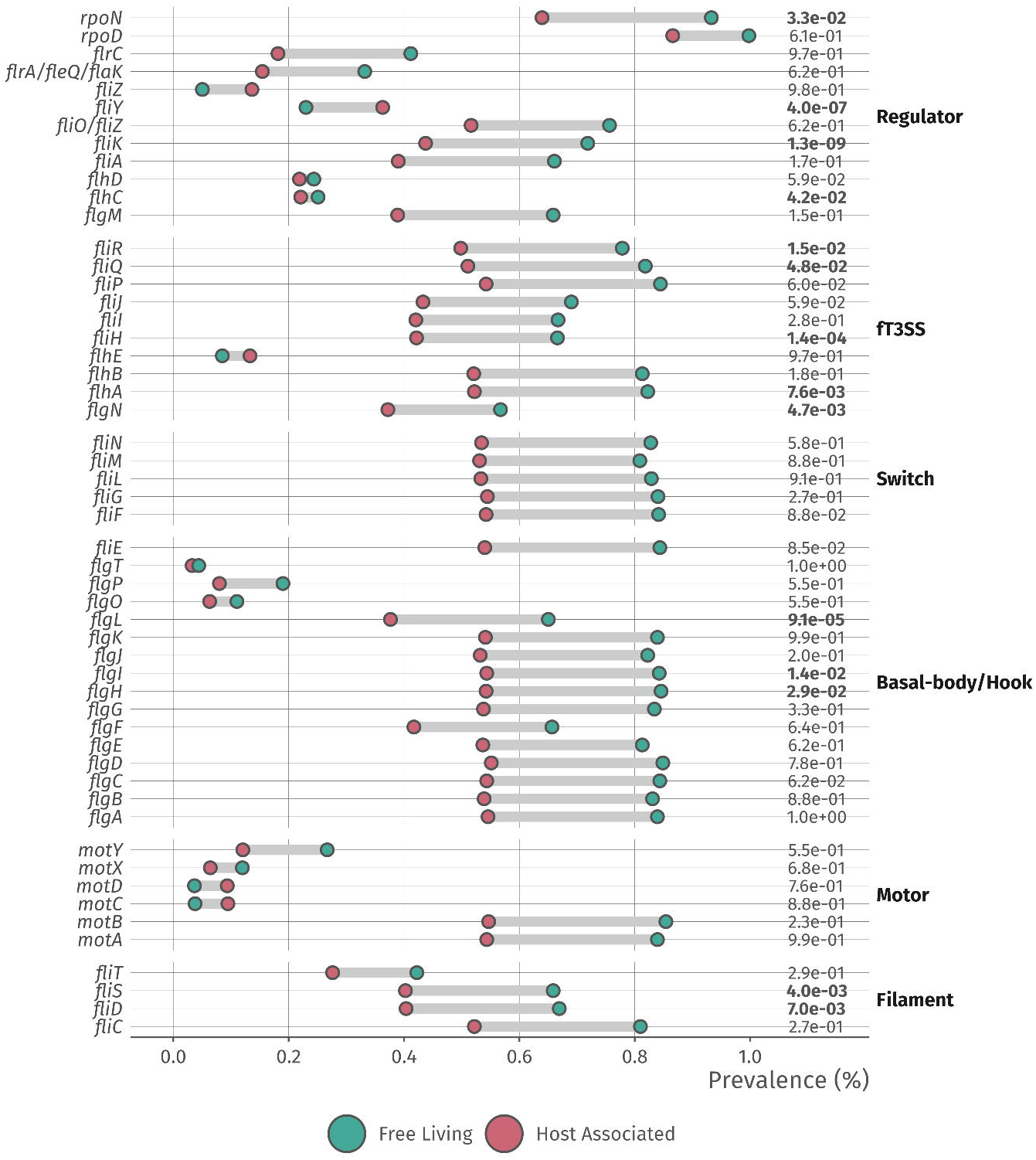
Detected prevalence of flagellar genes in host-associated and free-living bacteria. Observed prevalence of each flagellar gene in genomes of free-living and host-associated bacteria (including endosymbionts). The genes were detected by homology to the reference KEGG orthologs families. In almost all cases, flagellar genes are more prevalent in free-living bacteria. False discovery rates from the phylogenetic logistic regression on the right side of graph show the significance of the effects of host-association on the prevalence of each gene.

### Host-adaptation proceeds through neutral and diversifying selection

We investigated the effect of host-association on the degree of conservation of the detected flagellar gene sequences by phylogenetic linear regression. Host-association resulted in significantly lower conservation of 22 out of 53 flagellar genes (Fig. 2, Supplementary Data 5). However, the effect of host-association, on average, had a ten times smaller estimated impact than genome size on the sequence conservation. This suggests that, also in this case, genetic drift was the primary driver of sequence polymorphism differences. Host-association was linked to higher diversification of sigma-factor encoding genes (*rpoD, rpoN* and *fliA),* and of 5 genes of the fT3SS: *fliH*, *fliI*, *fliiJ*, *fliQ* and *flhB*. Other affected genes are the alternative master regulator *flrA*/*fleQ*/*flaK*, the regulator *flrC*, the switch complex genes *fliF* and *fliL*, the hook-filament junction gene *flgL*, the basal-body gene *flgP*, the filament cap gene *fliD*, the stator gene *motB* and the adaptor protein gene *fliE,* and the accessory motor genes *motXY.* Similarly to what we observed in the prevalence data, we did not find evidence that host-association lead to an increase in the polymorphism of *fliC*.

**Figure 2.**
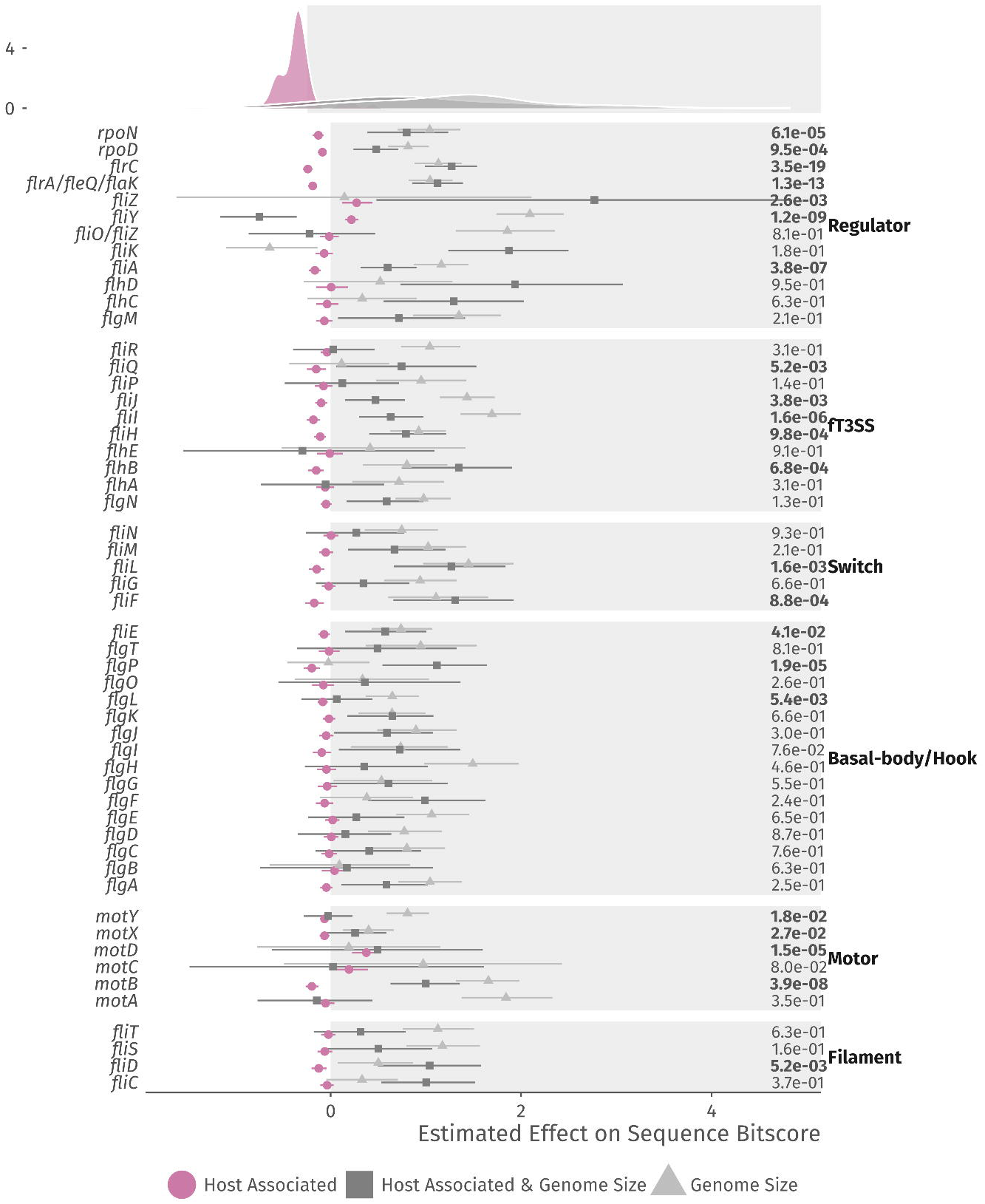
Estimates of host-association effects on flagellar gene conservation. Estimated effects and standard error of host-association, genome size and their interaction on the standardized bitscores of each gene, calculated using a phylogenetic linear regression. False discovery rates on the right side of graph show the significance of the effects of host-association. Density curves on top of the graph show the distribution of the effects for each term, highlighting the lower but consistently negative effects of host-association on sequence conservation.

## Discussion

For host-associated bacteria, loss of flagellar genes may help to evade immune detection while reallocating resources from motility to growth [10, 17]. To explore whether convergent adaptive dynamics influence the evolution of the flagellar assembly pathway in host-associated bacteria, we investigated the prevalence, redundancy, and conservation of 55 orthologs encoding flagellar regulators and building blocks, across a comprehensive genomic dataset of 1839 representative Pseudomonadota, encompassing both host-associated and free-living species.

Among bacteria harbouring *fliC*, encoding the essential filament protein, we identified three broad clusters of flagellar orthologs: rare, accessory, and core genes. Key regulatory genes (*flhC*, *flhD*, *flrA/fleQ/flaK*) exhibited low prevalence. Transcriptional regulators are known to be more evolutionary flexible than their targets [61], a condition that preserves the overall network structure even when the regulatory components themselves may vary [62]. For instance, previous studies have shown that *Pseudomonas fluorescens* can reintegrate flagellar functionality by recruiting a nitrogen uptake regulator following inactivation of *fleQ* [63]. A set of 23 orthologs, including components of the fT3SS, the switch complex, and parts of the basal body, hook, and stator, consistently co-occurred with *fliC*. The composition of this set of flagellar core genes aligns largely with prior studies [12].

Based on our comparative analysis we suggest that flagellar gene loss and diversification in host-associated bacteria are primarily driven by genetic drift and relaxed selection, with convergent positive selection acting only on a subset of genes. This is evidenced by the near-complete absence of flagellar genes in several exclusively host-associated orders from our genomic collection, such as Pasteurellales, Oceanospirillales, and Rickettsiales. Even in orders containing both host-associated and free-living strains, we found that flagellar gene prevalence is consistently lower in host-associated bacteria. Notable examples include Enterobacterales, Legionellales, Rhodospirillales, and Pseudomonadales. Interestingly, we also identified exceptions where free-living strains lack flagellar genes altogether (Moraxellales, Kangiellales, Acidiferrobacterales) or exhibit reduced prevalence compared to their host-associated relatives (Xanthomonadales, Aeromonadales). As our sample only included representative genomes, we cannot exclude that strain-level variability within these orders might produce some exceptions due to, for instance, horizontal gene transfer. Members of the order Moraxellales contain known opportunistic pathogens, and, at least in the case of the genus Psychrobacter, their evolution might have involved phases of host-association [64]. Several species of marine bacteria have experienced extensive losses of genetic material, although the drivers of reduction are likely different than those suggested for host-associated bacteria [65]. The Kangiellales are represented in our collection by 4 genomes of marine bacteria, with a genome size ranging from 2.4 to 2.8 Mb (for reference, *Escherichia coli* K12 has a genome of 4.6 Mb). Our models predict that both host-associated and free-living bacteria with smaller genomes are less likely to be flagellated. This does not only apply to marine bacteria, but also to [65] Acidiferrobacterales, which are autotrophic bacteria, able to oxidise sulphur and partly iron [66, 67]. Their ecological niche imposes a careful energetic budgeting and a predominantly non-motile lifestyle, as biofilms are fundamental for the leaching process [68]. Finally, the orders Xanthomonadales and Aeromonadales contain known plant and animal pathogenic and opportunistic species, for whom the evolution of flagellar genes is intertwined with the characteristic of their lifecycles. For instance, we detected no flagellar genes in the genome of *Xylella fastidiosa* Temecula1, which is an insect-borne plant pathogen. On the contrary, flagellar genes were commonly present in *Xanthomonas* spp., which is typically found in the phyllosphere, thus not transmitted by a symbiotic insect [69]. Flagellated host-associated bacteria that rely on motility for both survival outside the host and for host infection can adapt to evade the immune system [27, 70]. While some pathogenic species from the genus *Yersinia* have lost their flagellar genes, others regulate the expression of flagellar assembly in a temperature-dependent manner [71], enabling them to evade the immune system upon entering a mammalian host.

Our models link most differences in flagellar gene content to decreases in genome size, highlighting the role of genetic drift in the degeneration of the flagellar pathway. Importantly, while often considered neutral, genetic drift may include adaptive biases, as suggested by the positional and functional characteristics of lost genes [5]. Positive selection also played a significant, albeit smaller, role in shaping the flagellar regulon of host-associated bacteria, as evidenced by the estimated effects of host-association on both prevalence of flagellar genes and measures of their conservation.

We did not observe a significant effect of positive selection on prevalence and conservation of the filament protein *fliC* itself, aligning with prior studies suggesting its evolution is constrained by structural requirements and the high energetic costs of filament assembly [7]. Genes essential for filament elongation (*fliS*, *flhA*) were less prevalent in host-associated bacteria than in free-living bacteria, which could potentially lead to a reduction in immunogenicity of flagella of host-associated bacteria. Furthermore, we found a link between host association and lower copy numbers of commonly supernumerary flagellar genes (*motAB*, *fliL*, *fliC*), and increased diversification of a substantial fraction of flagellar orthologs. We found that host-associated bacteria disproportionately lose *flhC*, a mutation that would eliminate transcription of the entire flagellar regulon and might therefore predate the loss of *fliC*. They also exhibit diversification of *fliA*, which governs the expression of late-stage flagellar assembly genes and allows the fine-tuning of the motility phenotype, as suggested by a previous study on *Escherichia coli* [36]. In contrast, we did not observe direct effects of host-association on *flgM*, another regulatory gene proposed by the same study [36] to play a significant role in motility remodelling. Interestingly, *fliY* is one of the few genes to be more prevalent in host-associated than in free-living bacteria, which hints at an unknown role in the context of associations. FliY is a phosphatase that is part of the switch complex, which plays a role in the transduction of chemotactic signals [72], and its inactivation results in impaired motility and virulence [21].

This study has limitations. First, lifestyle classifications derived from isolation metadata are inherently uncertain, and annotations reflect only the conditions at the time of sampling rather than the full ecological breadth of each lineage. Although sensitivity analyses mitigate this concern, misclassification remains possible. Second, gene detection based on KEGG HMMs provides conservative presence/absence estimates and has limited sensitivity for poorly conserved flagellar components such as FliJ, FlgE, FliK, and FliD/ST. In addition, our pipeline does not incorporate synteny, gene-neighbourhood information, or remote homology searches, and therefore cannot distinguish complete flagellar systems from partial or degenerate remnants. Relatedly, polar and lateral flagellar systems are not resolved, and deep horizontal acquisition events cannot be identified with confidence. Finally, the evolutionary patterns reported here cannot fully separate genetic drift from relaxed selection, nor do they address potential neofunctionalization or adaptive diversification of retained genes. Future work combining targeted mutagenesis, experimental evolution, and structural or synteny-based annotation would help to refine and validate these inferences.

## Conclusions

In host-associated bacteria, reduced reliance on motility in nutrient-rich and relatively stable host environments, combined with demographic processes that facilitate the fixation of mutations under genetic drift and relaxed selection, likely contributes to the extensive loss of flagellar genes. At the same time, adaptation to specific host-associated niches may drive diversification within the remaining repertoire, allowing lineages to explore alternative functional trajectories. The patterns described in this study underscore the evolutionary flexibility of bacterial systems and provide a basis for future work examining the molecular and ecological forces that shape microbial adaptation within host environments.

## Resource Availability

### Data and code availability

Original code is publicly available at https://github.com/rsiani/phylogenomic_flagella as of the date of publication. Intermediate results are available at https://doi.org/10.5281/zenodo.17651454. Further information is available upon request from corresponding author, Roberto Siani.

## Acknowledgments

We thank Karin Pritsch for her contribution to revising the manuscript. This work was supported by the Deutsche Forschungsgemeinschaft (priority program SPP2125, project no. 401867691, awarded to Michael Schloter and Caroline Gutjahr).

## Author Contributions

R.S. designed the research study, conducted the analysis and wrote the manuscript; C.G. and M.S. contributed to the conceptualization of the study, supervised investigation and interpretation of results, edited and reviewed the manuscript.

## Supplemental Information

Data S1-S5

---

**Supplementary Figure 1.**
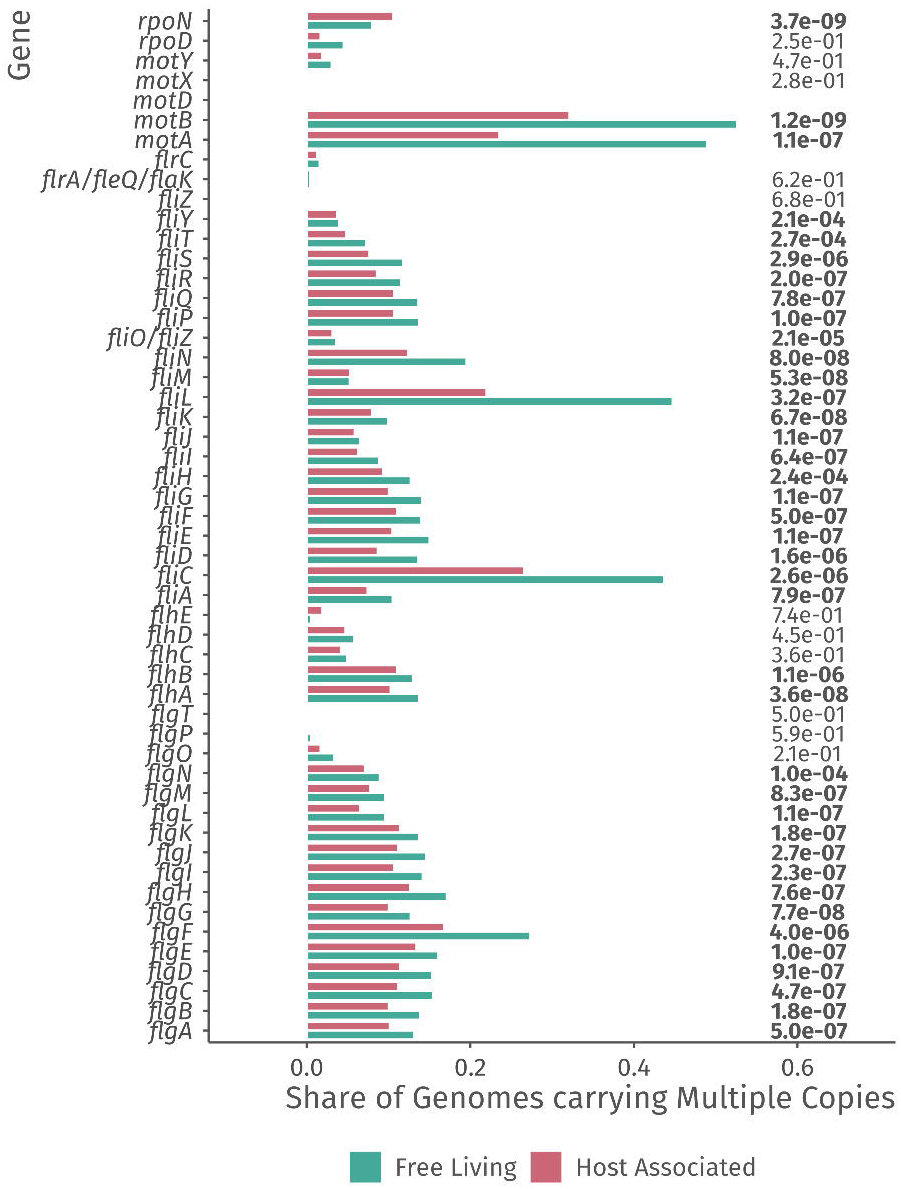
Host associated bacteria harbor less copies of supernumerary genes. Share of genomes, divided into the free-living and host-associated groups, carrying more than one copy of each gene. False discovery rates from the phylogenetic Poisson regression on the right side of graph show the significance of the effects of host-association on the number of copies of each gene. Multiple copies are consistently detected for the filament gene *fliC*, the motor genes *motA* and *motB,* and the switch gene *fliL,* with significant differences between free-living and host-associated bacteria.

